# Fiber Type and Stimulus Determine Progression of Skeletal Muscle Atrophy

**DOI:** 10.1101/2025.11.13.687882

**Authors:** Gregory L. Maas, Marcus P. Mullen, Brooke D. Shepard, Jack F. Gugel, Dakota R. Hunt, Virginia Ferguson, Sarah Calve, Thomas G. Martin, Leslie A. Leinwand

## Abstract

**Background:** Skeletal muscle atrophy is prevalent worldwide and is a major detractor from length and quality of life. It is often diagnosed and treated as a single disorder, but the causal stimuli and progression of atrophy vary widely. Malnutrition and disuse are two common causes of muscle atrophy, and despite their prevalence and extensive characterization, there have been no direct comparisons of how these two types of atrophy progress and whether they differentially affect skeletal muscle fiber types. The purpose of this study is to directly compare atrophy from fasting and disuse and provide a transcriptomic resource for future research on both conditions.

**Methods:** We fasted or hindlimb suspended (HS) two cohorts of 12-week-old female C57/bl6 mice. Mice were fasted for up to 72 hours to induce malnutrition atrophy or were hindlimb suspended for 0, 3, 7, 14, or 28 days to induce disuse atrophy. At each timepoint, mice were euthanized and three muscles (tibialis anterior (TA), extensor digitorum longus (EDL), and soleus) were weighed and collected for RNA sequencing. Atrophy progression and gene expression changes were compared across muscle fiber types and atrophy stimuli.

**Results:** We found differences in atrophy progression between muscle fiber types based on fiber twitch speed and atrophy stimulus. Fasted mice lost 25% of their body weight and 23% of fast-twitch TA mass with little change in soleus. In contrast, HS mice lost 40% of the slower-twitch soleus but the effect on the TA was negligible. Gene expression varied in response to both atrophy stimuli, but a greater number of genes changed with fasting compared to HS in the EDL and soleus. By muscle type, a greater transcriptional shift occurred in the EDL with fasting while the soleus showed more gene changes during HS. Enrichment analysis of transcriptional changes showed similarities (downregulation in muscle growth pathways) and differences (increased fatty acid metabolism in fasting and increased neuronal activity in HS) between atrophy stimuli.

**Conclusions:** Atrophy progression varies based on stimuli and muscle fiber type. This study provides a large, matched data set where the effects of different atrophic stimuli can be easily and directly compared in multiple fiber types. To our knowledge, this is the first study to closely compare these two atrophy stimuli in a muscle type-specific context. This work demonstrates that atrophy is not a single disorder and that the development of therapies may need to be tailored to the atrophic stimulus.

## Introduction

Maintaining muscle mass and function are fundamental to improving quality of life and extending healthspan [1-2]. Skeletal muscle atrophy, the progressive deterioration of muscle mass and function, is increasingly prevalent as global populations age and experience compounding lifestyle factors [3-5]. Atrophy accelerates with age and is worsened by inactivity and malnutrition, or diseases including cancer, muscular dystrophy, and immune deficiencies [6-7]. For example, sarcopenia, the loss of muscle mass and function with aging, typically begins around 35 years of age with a loss of ∼1% of muscle mass per year that increases to ∼3% or more lost per year by age 60 [8]. Lifestyles in developed countries are increasingly sedentary, decreasing the amount of physical activity that adults regularly engage in and raising their risk for muscle atrophy [9]. Compounding this is a global crisis of malnutrition; nearly 10% of the world is malnourished [10-11]. Both inactivity and malnutrition contribute to poor muscle growth and maintenance [12-13]. Despite its prevalence and inevitability with age, there are no effective pharmaceutical treatments for muscle atrophy [14]. Atrophy can be mitigated through lifestyle changes such as proper nutrition and exercise, but such general preventative measures are not accessible for all people. Thus, there is a need for therapeutic modalities that target atrophy at the molecular source.

The base cause of muscle atrophy is consistent across stimuli and conditions. Skeletal muscle cells (myocytes) are long-lived cells that rely on continual protein turnover to maintain proper function [15]. Damaged proteins are degraded and replaced by newly synthesized ones, and even functional proteins may be degraded to be used as fuel [16-17]. In a homeostatic system, the rate of protein synthesis and degradation are equal, resulting in no change in net muscle protein [18]. In a muscle undergoing atrophy, the rate of protein degradation outweighs synthesis, causing a loss in protein which reduces muscle mass and impairs function [19]. While the end result and root cause of muscle atrophy between disuse and malnutrition are similar, the intermediary pathways can differ dramatically [20]. Muscle atrophy is not a single pathological process that can be defined by a single cellular signaling pathway.

In addition to proceeding via distinct intermediary pathways, malnutrition- and disuse-induced atrophy also have different effects on muscle sub-types [19, 20]. Malnutrition-induced atrophy preferentially degrades fast-twitch muscles, whereas unloading-induced atrophy has a stronger effect on slow twitch muscles [13, 21]. These differential susceptibilities likely reflect fiber type-specific differences in metabolism, protein turnover, and mechanical load dependence. While previous studies characterize these muscle groups and atrophy types individually, it is not clear how atrophy progresses and differs between disuse and fasting in a muscle type-specific context. An improved understanding of individual causes and specific progressions of atrophy will provide mechanistic insights and new treatment options for those affected by each type of atrophy rather than treating them analogously.

In this study, we sought to identify the molecular underpinnings of skeletal muscle atrophy induced by disuse versus fasting in a muscle-specific context. To explore this, we used two mouse models of skeletal muscle atrophy: hindlimb suspension (HS) and fasting. We performed bulk RNA-seq analysis of both fast- (tibialis anterior (TA) and extensor digitorum longus (EDL)) and slow-twitch (soleus) muscle fibers. We found that both the atrophy stimulus and muscle fiber type determined the severity and progression of muscle loss. Fasting affected both muscle types, but mass loss was greater in the fast-twitch TA and EDL. HS had minimal effects in the TA and EDL but greatly impacted the slow-twitch soleus. This study provides a resource for comparison of atrophy modalities across different skeletal muscle fiber types.

## Results

### Total body weight is reduced by fasting but not HS

To investigate how different atrophy stimuli affect body composition, we measured body and individual muscle weights of mice that were either fasted for 72 hours or were hindlimb suspended for 0, 3, 7, 14, or 28 days. Expectedly, fasted mice lost 25% of their total body mass (∼5g) after 72 hours (**Fig 1a**), with a daily loss of 1.8 – 4.8% body mass (∼1.5 g) per day (**Fig 1b**). Weight loss occurred rapidly early in the fast (5% daily), then slowed as fasting continued (1.8% daily). Interestingly, weight loss in fasting mice was driven by both lean muscle (3.1 g total average loss, p=0.0001) and fat mass (2.8 g total average loss, p=0.0004) (**Fig 1c**). HS mice, however, experienced a transient decrease in body weight immediately following suspension that was recovered at 28 days (**Fig 1d,e**), which has been previously reported after HS [22].

**Fig. 1.**
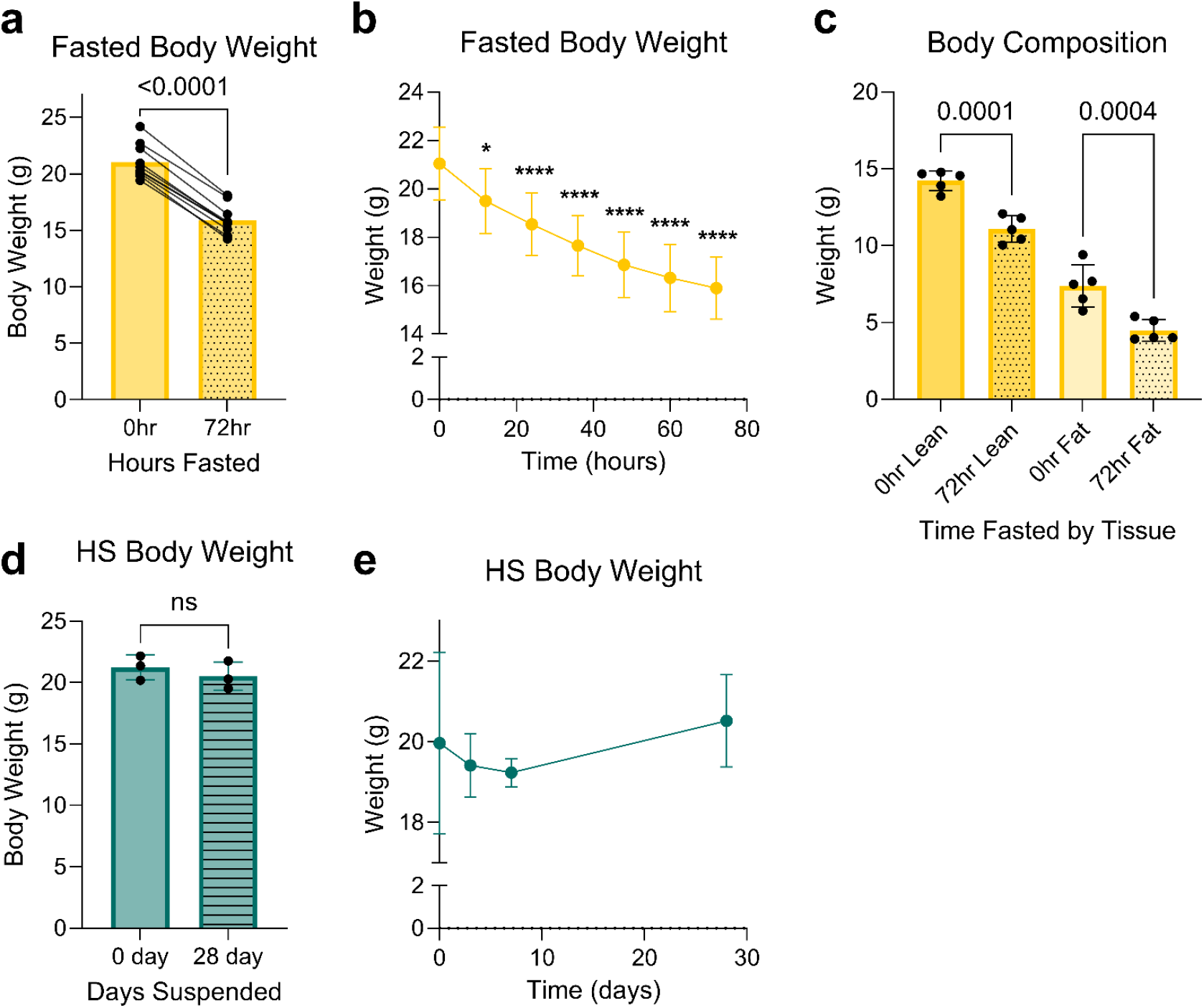
Fasting exerts a greater effect on total body weight than hindlimb suspension. **(a)** Paired mouse total body weight before and after 72 hours of fasting (n=10). **(b)** Decline in mouse body weights at 12-hour intervals after fasting onset (n=10). **(c)** Lean and fat mass of mice before and after 72 hours of fasting measured by DEXA Scan (n=5). **(d)** Mouse body weights before and after 28 days of HS (n=3). **(e)** Average mouse body weights at 0, 3, 7, and 28 days of HS (n=3). For a and d, data were analyzed by t-test. For b, c, and e, the data were analyzed by one-way ANOVA. When a significant interaction was identified, Tukey’s post-hoc test for multiple independent pairwise comparisons was used; *p < 0.05, ****p < 0.0001, ns = not significant.

### Atrophy progression with fasting and HS is dependent on muscle type

To measure the effects of fasting and disuse on different muscle types, TA, EDL, and soleus were collected from mice at each timepoint. TA mass decreased significantly during fasting but was unaffected by HS (**Fig 2a**). EDL mass decreased with fasting but was unchanged in HS (**Fig 2b**). Fasting induced a small, but significant loss in soleus mass while HS loss progressed from a tibia-normalized mass of 0.65 mg/mm at baseline to 0.22 mg/mm at 14 days suspended (**Fig 2c**). To explore potential differences in the underlying molecular regulation of atrophy in these two conditions, we next characterized global gene expression in each atrophy stimulus and muscle type to define transcriptome differences.

**Fig. 2.**
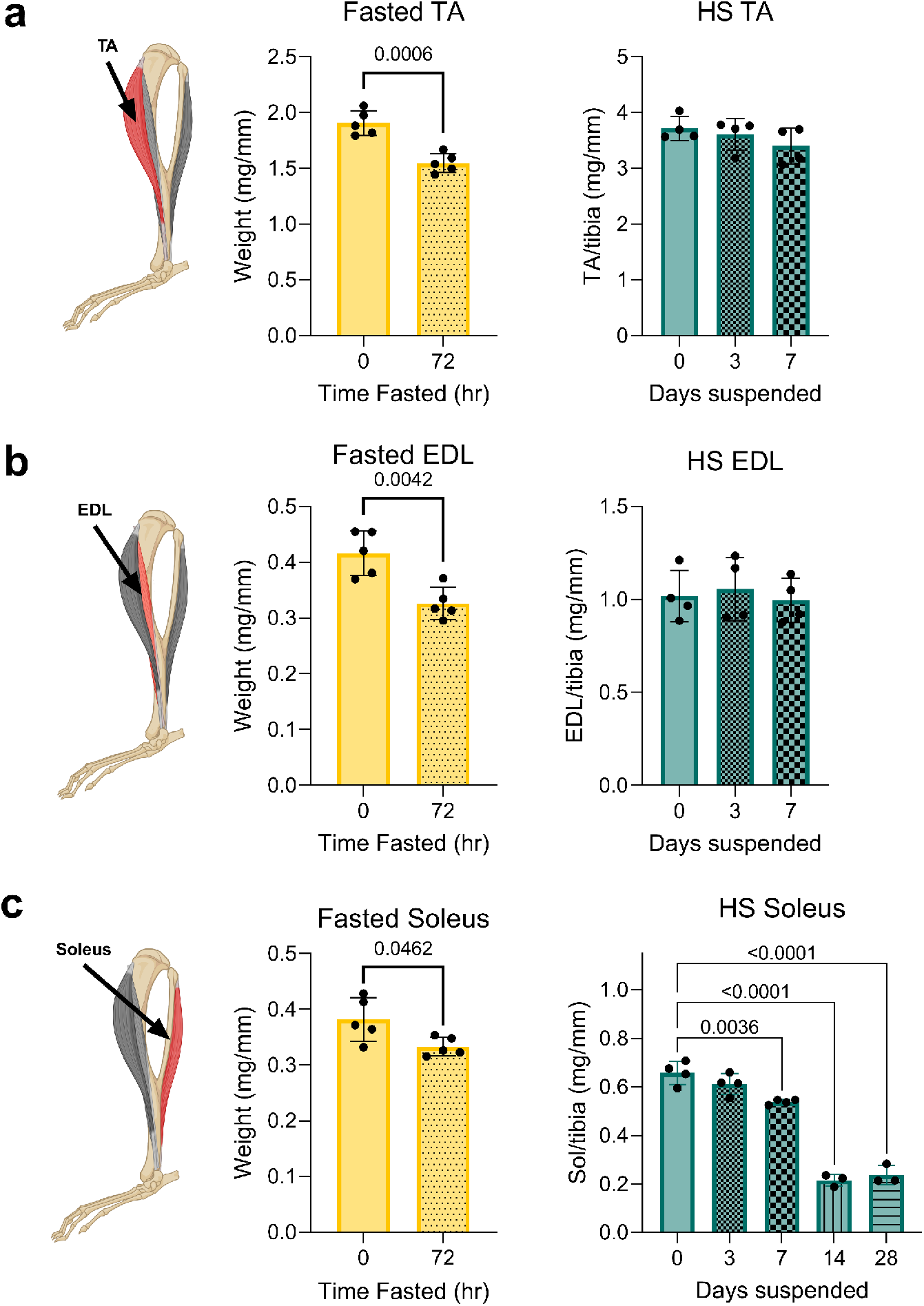
Slow- and fast-twitch muscles are differentially affected by fasting and hindlimb suspension. **(a)** Tibialis Anterior (TA) diagram (TA in red) and muscle weights normalized to tibia length after fasting (yellow, n=5) and HS (blue, n=4). **(b)** Extensor Digitorum Longus (EDL) diagram (EDL in red) and muscle weights normalized to tibia length after fasting (yellow, n=5) and HS (blue, n=4). **(c)** Soleus diagram (soleus in red) and muscle weights normalized to tibia length after fasting (yellow, n=5) and HS (blue, n=4). Fasting analyzed with Welch’s t-test. HS analyzed by one-way ANOVA.

### Gene expression changes are dependent on both the atrophy stimulus and the muscle fiber type

Our lab has previously demonstrated that atrophy progression depends on stimulus and muscle-type differences in protein turnover [23]. Therefore, we investigated the transcriptional differences caused by fasting and disuse in the context of slow- (soleus) and fast- (EDL) twitch muscle by bulk mRNA sequencing (**Fig S1**). Muscles were collected after mice were exposed to 3 days of fasting or HS. In the EDL, we found genes relating to growth and cell fate (*Foxo6os* and *Nrep*) decreased with fasting, while pathways related to fatty acid metabolism (*Acot1* and *Acss1*) increased (**Fig 3a**). The predominant change induced by HS in the EDL was a suppression of the immune response (e.g., *Rcan1*) (**Fig 3a**). Strikingly, the number of genes changing in the EDL was different based on the atrophy stimulus, with 3,447 genes being unique to fasting, 141 genes being unique to suspension, and 171 genes affected by both stimuli (**Fig 3a**). In the soleus, genes with a role in hypertrophy and cell proliferation (e.g., *Foxo6os, Nrep*, and *Myoz3*) decreased with fasting, while genes associated with glycolysis increased (e.g., *Acot1*) (**Fig 3b**). In the soleus, 2,187 genes were uniquely affected by fasting, 704 with suspension, and 368 genes whose expression was altered in both conditions (**Fig 3b**).

**Fig. 3.**
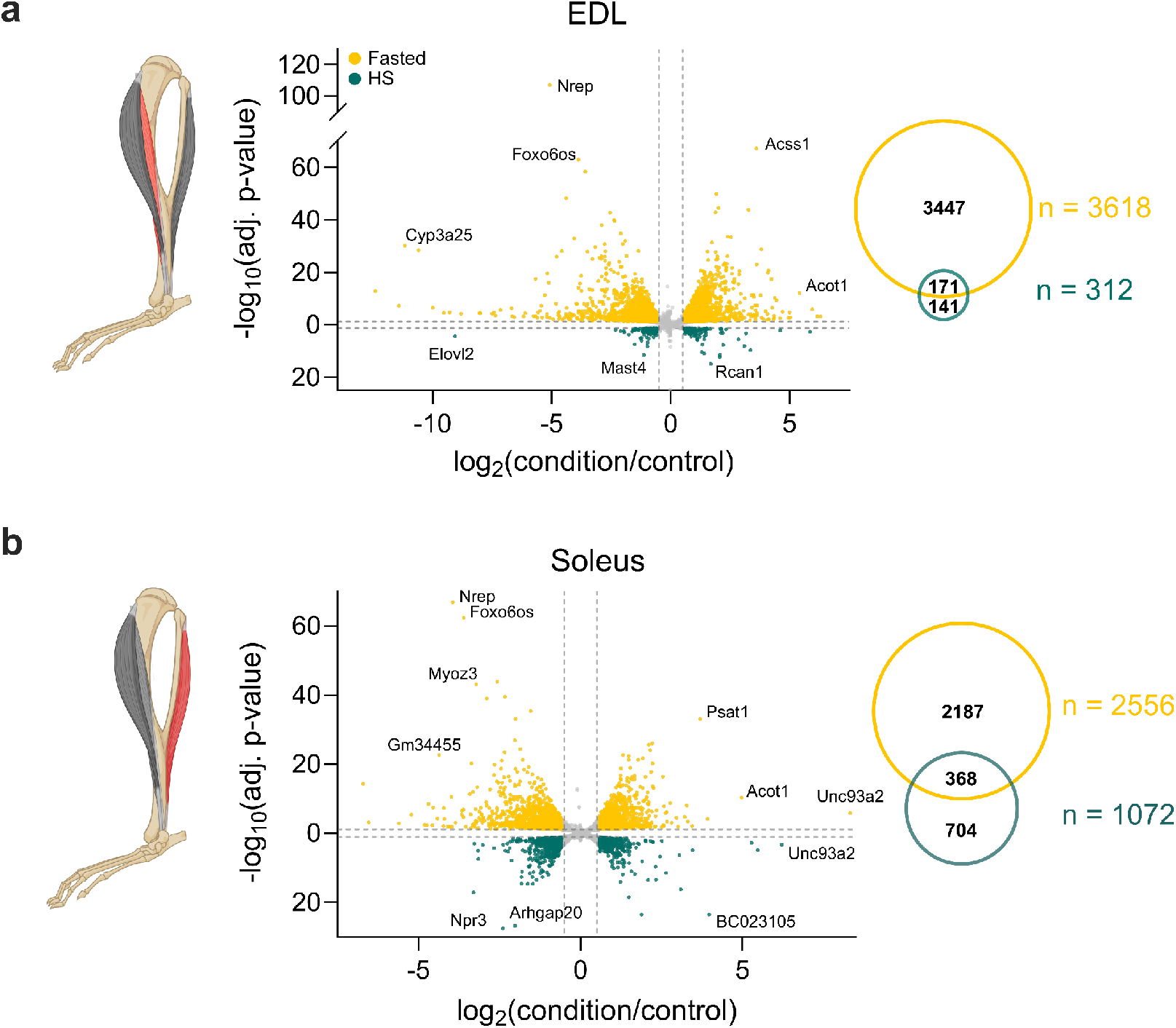
EDL and soleus have distinct transcriptional responses to atrophy stimuli. **(a-b)** Volcano plots depicting differential gene expression for Fast vs. Fed control (top, yellow) and Suspension vs. Unsuspended control (bottom, teal) in EDL (a) and soleus (b) alongside Venn diagrams representing unique and overlapping differentially expressed genes in each condition and muscle-type. Significance cutoff: adj. p value < 0.05; fold change cutoff: |fold change| > 1.5

### Some genes have dramatically different responses to atrophy stimuli in different muscle fiber types

To determine how the same gene changed in each atrophy stimulus between the soleus and the EDL, we graphed the fold change of each gene in the soleus versus its change in the EDL. Along with bioinformatic tools (see methods), this allowed us to directly compare which genes had a significantly different response to the atrophy stimuli in each muscle type. Gene expression is distinguished by being differentially increased in either the EDL or soleus. In Fasting, expression of 138 genes was higher in the EDL while 133 genes had higher expression in the soleus (**Fig 4a**). Enrichment analysis (GSEA) showed that genes related to oxidative energy production were elevated in the EDL after fasting (**Fig 4b**). Meanwhile in HS, 188 genes were more upregulated in the EDL compared to 138 that were higher in the soleus (**Fig 4c**). Enrichment analysis after HS identified that genes related to amino acid catabolism and metabolism were increased in soleus (**Fig 4d**). Considering both our body and muscle weight measurements and transcriptomic analyses, these data reveal that fasting-induced atrophy exhibits a greater whole-body mass loss with acute atrophic effects on fast twitch muscles, while HS-induced atrophy is less severe globally but greatly impacts the mass and transcriptional profile of the slow-twitch soleus muscle.

**Fig. 4.**
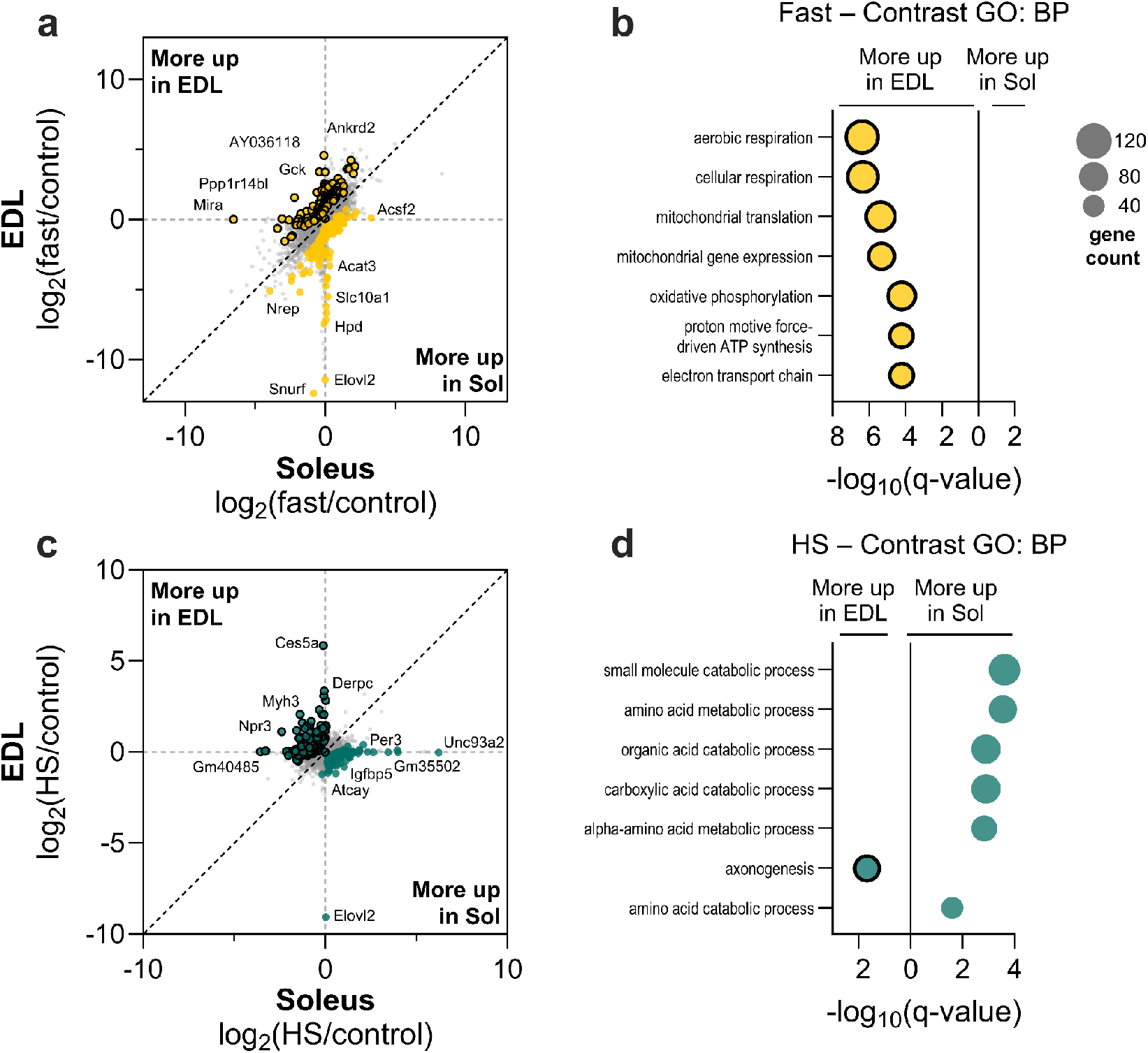
Transcriptional responses to disuse and nutrient depravation depend on muscle group. **(a)** Paired muscle-type–specific fold-change plots highlighting genes whose response to fasting differed significantly (adj. p value < 0.05) between EDL and soleus; dotted line = line of identity. **(b)** Top seven most significant terms from Gene Ontology (GO) Biological Process GSEA of genes whose expression differed between soleus and EDL in fasting **(c)** Paired muscle-type-specific fold-change plots highlighting genes whose response to suspension differed significantly (adj. p value < 0.05) between EDL and soleus; dotted line = line of identity. **(d)** Top seven most significant terms from Gene Ontology (GO) Biological Process GSEA of genes whose expression differed between soleus and EDL in suspension. Genes more up in the EDL are outlined in black; gene count = core enrichment genes

## Discussion

Muscle atrophy severely impacts quality of life, reducing mobility, strength, endurance, and quality of life [6-8]. With its prevalence across populations [3] and adverse effects on quality of life, it is important to understand and mitigate atrophy to increase healthspan. Despite extensive study, muscle atrophy is often viewed as a single condition with a single prognosis and progression. In truth, muscle atrophy has myriad causes; here we characterize muscle atrophy in the two most common causes, malnutrition (via fasting) and disuse (via hindlimb suspension), showing that atrophy from these stimuli differs greatly in progression and has distinct effects on muscle fiber types.

Malnutrition deprives the muscle of new protein [24]. Existing proteins are subsequently proteolyzed due to the lack of new dietary proteins, thereby reducing body and muscle mass. Malnutrition affects all muscles, but the resulting atrophy is especially acute in fast-twitch muscle fibers that are highly energy-intensive and are activated only when needed [21, 23]. In this study, the fast-twitch muscles (TA and EDL) lost ∼20% of their starting weight, while the soleus, a slow-twitch muscle, lost only 12% after 72 hours of fasting. All three muscles lost a statistically significant amount of mass, exhibiting fasting-induced atrophy in all muscle types studied. However, the EDL lost significantly more mass from fasting than the soleus. This preferential loss in fast-twitch muscles during fasting is due, in part, to the role of fast-twitch muscle as a glycogen reservoir [26]. As fasting reduces intramuscular glucose availability, fast-twitch muscle must catabolize its own carbohydrate stores and eventually its own protein, reducing muscle size and function. These freed sugars and amino acids can then enter glycolysis and the TCA cycle, sustaining essential physiological processes as nutrient deprivation continues [25-27]. As energy reservoirs, fast-twitch muscles are highly susceptible to nutrient deprivation. Notably, we found the degree of muscle catabolism and effect on fatty acid metabolism is greater in the EDL than the soleus. Further, the biological processes downregulated in fasting differ between fast- and slow-twitch muscles. The top ten downregulated GO terms in the soleus implicate oxidative phosphorylation and the electron transport chain, suggesting a switch in energy production pushing the usually oxidative soleus more glycolytic as fasting progresses.

Disuse removes strain from postural muscles which, through mechanosensing pathways, results in muscle degradation [28-29]. The effects of unloading were mostly confined to the soleus, which decreased in mass by 20%, while TA mass decreased by just 9% after 7 days of suspension. Gene expression changes in HS were also more pronounced in the soleus, where three times as many genes (1,072 versus 312) were differentially expressed compared to the EDL. The soleus is a slow-twitch oxidative muscle which preferentially used aerobic metabolism to generate ATP [25]. We observed significant activation of pathways involved in synaptic protein regulation in both muscle types, but this was much higher in the soleus, possibly indicating a mechanosensing response to muscle unloading. As a postural muscle, the soleus is almost always under strain [32]. Removing load results in a disuse signal starting with dephosphorylation of AMPK and cascading to cause degradation of the unused muscle [29, 31].

Atrophy stimuli and their associated signaling pathways have been well studied individually; however, to our knowledge, there are no examples of studies where multiple stimuli and muscle types are compared directly. For example, molecular pathways mediated by AMPK, FoxO transcription factors, NF-ĸB, and myostatin regulate atrophy, but are activated to different extents in each atrophy condition [32-34]. Nutrient availability, energy consumption, and level of use all influence how these pathways interact and trigger degradation in each muscle type. However, the precise interplay of each of these pathways between atrophy stimuli is still poorly understood. The power of this study lies in the direct matched animal comparison of the two atrophy conditions while accounting for fiber type. By using age-, sex-, and weight-matched mice, we can readily compare mass loss and transcriptomic changes in TA, EDL, and soleus between atrophy types. Excitingly, this direct comparison allows for the discovery of heretofore overlooked genes. For example, the long noncoding RNA *Foxo6os* was significantly downregulated in both EDL and soleus during fasting and its overexpression has recently been implicated in cardiac hypertrophy [37]. Further investigation into this lncRNA could produce novel therapeutic avenues. The parsing of disuse and malnutrition induced atrophy can inform individualized treatment strategies for each condition, but it also enables a more tailored treatment for those experiencing both stimuli simultaneously. For example, a patient confined to bed rest on a liquid diet could be treated for both forms of atrophy simultaneously, ideally with a single drug or cocktail that addresses both atrophy stimuli.

Different populations experience different atrophy stimuli, making it important to distinguish the cause of atrophy and understand its specific progression to better mitigate muscle loss and weakness. A one treatment-fits-all approach to muscle atrophy has limited efficacy for those most impacted by this disease. Our work provides a resource for two common stimuli of muscle atrophy. A deeper investigation into specific genes identified herein could provide insight into possible therapeutic avenues for atrophy.

Future studies examining muscular atrophy should be aware of these multilevel differences in atrophy stimulus and utilize this transcriptomic database when designing subsequent experiments to create thorough and holistic answers to muscle atrophy from different stimuli.

## Methods

### Mouse Husbandry

All mouse experiments were performed according to procedures approved by the IACUC of University of Colorado Boulder.

### Fasting Paradigm

Mice were maintained in 12-hour light-dark cycles at 22°C with 50% relative humidity. Mice were fed with a standard rodent chow diet (18% protein and 6% fat; Envigo Teklad 2018). C57BL/6J (Cat. 000664) mice were obtained from The Jackson Laboratory.

Mice were divided into two groups: Fed Control and Fasted. The fed group received food *ad libitum* while the fasted group had their food removed and were transferred to a clean cage where they were fasted for up to 72 hours. Mice were euthanized via midline thoracotomy followed by rapid excision of the heart under deep isoflurane-induced anesthesia (3% isoflurane delivered at a rate of 0.8 liter/minute). Sufficient anesthetic depth was confirmed by lack of response to physical stimuli. Whole body weight was measured and blood was collected via the posterior vena cava, allowed to clot, and spun for 10 minutes at 3,000 rpm to isolate serum, then frozen at -80 °C. Tibialis anterior (TA), extensor digitorum longus (EDL), and soleus muscles were dissected, weighed, and flash frozen to store at -80 °C. Muscle weights were normalized to tibia length. The protocol for this study was approved by the Institutional Animal Care and Use Committee of the University of Colorado at Boulder.

### Hindlimb Suspension

11-week-old, female, wild type C57BL/6 mice arrived from Jackson labs and were allowed to acclimate to our facility for 7 days before the start of the experiment (12-week-old upon experiment start). Upon arrival, mice were housed in standard polycarbonate cages with 7090A Teklad Aspen sani-chip bedding, were maintained on a 12:12 h light-dark cycle, and allowed ad libitum access to water and standard rodent diet (2918 Teklad).

Once acclimated, mice were randomly assigned to the control group or suspension group. Mice underwent HS for 0, 3, 7, 14, or 28 days (n = 4) using custom suspension cages that included a large hamster cage as a base (Allentown Cages, Allentown, NJ), and two custom, laser-cut parts: a cage topper to allow for filtered air flow and water bottles, and flat mesh to allow mice to grip the cage bottom without slipping. The tails of suspended mice were attached with low-irritation medical tape to a small swivel device made of a dowel and small hook. The swivel device connected to a wire that ran the length of the cage, allowing free rotation and mobility across the entire cage. Mice were hindlimb suspended at a 30-degree head down-tilt [S1, S2] and socially housed with one suspended mouse and one control (unsuspended) mouse to reduce stress [S3]. To confirm that suspended mice were not standing on control mice to avoid unloading, we reviewed 100 hours of video footage with this setup and did not observe any loading of the limbs due to social housing. To avoid further unnecessary loading of the hindlimbs, we limited bedding and enrichment to ensure mice did not stand on piles of cage materials. Mice were health checked twice daily to ensure mice did not slip out of swivels, slip down farther than 30 degrees incident to the cage base, or experience adverse health effects.

Mice were euthanized via CO_2_ asphyxiation and cervical dislocation. Whole body weight was measured and blood was collected via the posterior vena cava, allowed to clot, and spun for 10 minutes at 3,000 rpm to isolate serum, then frozen at -80 °C. Tibialis anterior (TA), extensor digitorum longus (EDL), and soleus muscles were dissected, weighed, and flash frozen to store at -80 °C. Muscle weights were normalized to tibia length. The protocol for this study was approved by the Institutional Animal Care and Use Committee of the University of Colorado at Boulder.

### DXA Scan

At treatment endpoint, mice were euthanized by CO_2_ then confirmed by cervical dislocation before being transported to University of Colorado Anshutz Medical Campus for DXA scan. Scans were completed on a DXA Body Composition Analyzer, Model InAlyzer manufactured by MEDIKORS Inc. Mice were individually arranged on the scanning area and exposed to low-dose x-ray. Radiation absorbance was measured and recorded to report lean, fat, and bone content of each mouse.

### RNA extraction

RNA was extracted from tissue via TRIzol/chloroform extraction and was purified using RNeasy Minikit (Qiagen). On column DNase digest was performed to remove any genomic DNA contamination. Samples were then sent to Novagene for RNAseq.

### RNA sequencing and Enrichment Analysis

Bulk RNA from mouse EDL and soleus (n = 2 per condition) was sent to Novogene Corporation (Sacramento, CA) for poly-A enrichment and short-read mRNA sequencing at a minimum 50 million read-pair depth per sample. For analysis, Nextflow version 21.10.6 was used to run the nf-core/rnaseq pipeline version 3.9.0 [S4, S5]. Gene count normalization and differential gene expression analysis was conducted with DESeq2 version 1.42.1 [S6]. For all designs, shrinkage was performed using apeglm [S7] to improve the visualization of genes. For comparisons of a given muscle between a condition (either fasting or suspension) and control group, genes were considered differentially expressed with a shrunken |fold-change| > 1.5 and with an FDR-adjusted p-value < 0.05. To gauge overall effects of fasting or suspension, we used the following design: muscle type + condition. To identify gene expression that responds differently in soleus and EDL after a given condition (either fasting or suspension), we added an interaction term: muscle_type + condition + muscletype:condition. Genes with an FDR-adjusted p-value < 0.05 were considered differentially expressed depending on muscle-type, and subsequent enrichment analysis was performed using the DESeq2 results. For example, this can mean the gene is upregulated in muscle one and down in muscle two, up in muscle one and unchanged in muscle two, or unchanged in muscle one and down regulated in muscle two. Any of these responses show differential gene expression between muscle types. All enrichment analyses were performed with GSEA [S8, S9] version 4.10.1, using gene sets ranked by the negative log2 FDR-adjusted p-value multiplied by the sign of the fold change. Gene sets with FDR-adjusted q-values < 0.05 were considered significant.

## Acknowledgements

We thank Ginger Johnson and Margeurite Kehler at the University of Colorado Anschutz Medical Center for access and assistance in acquiring all DXA data. We thank all OAR staff at University of Colorado Boulder for animal and veterinary care, especially Aaron T. Rothchild, Lindsay Larson, and Emily Bourgeois. Funding was provided by the National Institutes of Health (R01GM029090 to LAL, T32-GM142607 to MPM).

## References

[1] P. C. Guest, “New Therapeutic Approaches and Biomarkers for Increased Healthspan,” in Reviews on New Drug Targets in Age-Related Disorders: Part II, P. C. Guest, Ed., Cham: Springer International Publishing, 2021, pp. 1–13. doi: 10.1007/978-3-030-55035-6_1.

[2] Y. Guan and Z. Yan, “Molecular Mechanisms of Exercise and Healthspan,” Cells, vol. 11, no. 5, p. 872, Jan. 2022, doi: 10.3390/cells11050872.

[3] F. Petermann-Rocha et al., “Global prevalence of sarcopenia and severe sarcopenia: a systematic review and meta-analysis,” J. Cachexia Sarcopenia Muscle, vol. 13, no. 1, pp. 86–99, 2022, doi: 10.1002/jcsm.12783.

[4] R.-Y. Wu, W.-H. Sung, H.-C. Cheng, and H.-J. Yeh, “Investigating the rate of skeletal muscle atrophy in men and women in the intensive care unit: a prospective observational study,” Sci. Rep., vol. 12, p. 16629, Oct. 2022, doi: 10.1038/s41598-022-21052-3.

[5] L. Jun, M. Robinson, T. Geetha, T. L. Broderick, and J. R. Babu, “Prevalence and Mechanisms of Skeletal Muscle Atrophy in Metabolic Conditions,” Int. J. Mol. Sci., vol. 24, no. 3, p. 2973, Jan. 2023, doi: 10.3390/ijms24032973.

[6] D. Scott, L. Blizzard, J. Fell, and G. Jones, “The epidemiology of sarcopenia in community living older adults: what role does lifestyle play?,” J. Cachexia Sarcopenia Muscle, vol. 2, no. 3, pp. 125–134, Sept. 2011, doi: 10.1007/s13539-011-0036-4.

[7] O. Rom, S. Kaisari, D. Aizenbud, and A. Z. Reznick, “Lifestyle and Sarcopenia—Etiology, Prevention, and Treatment,” Rambam Maimonides Med. J., vol. 3, no. 4, p. e0024, Oct. 2012, doi: 10.5041/RMMJ.10091.

[8] A. J. Cruz-Jentoft et al., “Sarcopenia: European consensus on definition and diagnosis,” Age Ageing, vol. 39, no. 4, pp. 412–423, July 2010, doi: 10.1093/ageing/afq034.

[9] T. Strain et al., “National, regional, and global trends in insufficient physical activity among adults from 2000 to 2022: a pooled analysis of 507 population-based surveys with 5·7 million participants,” Lancet Glob. Health, vol. 12, no. 8, pp. e1232–e1243, Aug. 2024, doi: 10.1016/S2214-109X(24)00150-5.

[10] “Fact sheets - Malnutrition.” Accessed: Oct. 14, 2025. [Online]. Available: https://www.who.int/news-room/fact-sheets/detail/malnutrition

[11] “Malnutrition: share of people by region 2023,” Statista. Accessed: Oct. 14, 2025. [Online]. Available: https://www.statista.com/statistics/273291/number-of-people-with-malnutrition-worldwide/

[12] J. Lopes, D. Russell, J. Whitwell, and K. Jeejeebhoy, “Skeletal muscle function in malnutrition,” Am. J. Clin. Nutr., vol. 36, no. 4, pp. 602–610, Oct. 1982, doi: 10.1093/ajcn/36.4.602.

[13] A. J. Sargeant, C. T. M. Davies, R. H. T. Edwards, C. Maunder, and A. Young, “Functional and Structural Changes after Disuse of Human Muscle,” Clin. Sci., vol. 52, no. 4, pp. 337–342, Apr. 1977, doi: 10.1042/cs0520337.

[14] G. Bahat and S. Ozkok, “The Current Landscape of Pharmacotherapies for Sarcopenia,” Drugs Aging, vol. 41, no. 2, pp. 83–112, Feb. 2024, doi: 10.1007/s40266-023-01093-7.

[15] D. J. Millward, P. J. Garlick, D. O. Nnanyelugo, and J. C. Waterlow, “The relative importance of muscle protein synthesis and breakdown in the regulation of muscle mass,” Biochem. J., vol. 156, no. 1, pp. 185–188, Apr. 1976, doi: 10.1042/bj1560185.

[16] A. L. Goldberg, “Protein degradation and protection against misfolded or damaged proteins,” Nature, vol. 426, no. 6968, pp. 895–899, Dec. 2003, doi: 10.1038/nature02263.

[17] P. de Lange, M. Moreno, E. Silvestri, A. Lombardi, F. Goglia, and A. Lanni, “Fuel economy in food-deprived skeletal muscle: signaling pathways and regulatory mechanisms,” FASEB J., vol. 21, no. 13, pp. 3431–3441, 2007, doi: 10.1096/fj.07-8527rev.

[18] I. V. Hinkson and J. E. Elias, “The dynamic state of protein turnover: It’s about time,” Trends Cell Biol., vol. 21, no. 5, pp. 293–303, May 2011, doi: 10.1016/j.tcb.2011.02.002.

[19] P. Bonaldo and M. Sandri, “Cellular and molecular mechanisms of muscle atrophy,” Dis. Model. Mech., vol. 6, no. 1, pp. 25–39, Jan. 2013, doi: 10.1242/dmm.010389.

[20] Y. Wang and J. E. Pessin, “Mechanisms for fiber-type specificity of skeletal muscle atrophy,” Curr. Opin. Clin. Nutr. Metab. Care, vol. 16, no. 3, pp. 243–250, May 2013, doi: 10.1097/MCO.0b013e328360272d.

[21] T. Ogata, Y. Oishi, M. Higuchi, and I. Muraoka, “Fasting-related autophagic response in slow- and fast-twitch skeletal muscle,” Biochem. Biophys. Res. Commun., vol. 394, no. 1, pp. 136–140, Mar. 2010, doi: 10.1016/j.bbrc.2010.02.130.

[22] M. E. Rosa-Caldwell et al., “Female mice may have exacerbated catabolic signalling response compared to male mice during development and progression of disuse atrophy,” J. Cachexia Sarcopenia Muscle, vol. 12, no. 3, pp. 717–730, 2021, doi: 10.1002/jcsm.12693.

[23] J. Gugel, “Muscle Protein Turnover in Health and Disease,” Ph.D., University of Colorado at Boulder, United States -- Colorado, 2025. Accessed: Oct. 27, 2025. [Online]. Available: https://www.proquest.com/docview/3244752227/abstract/67F5510DAF7F4C3CPQ/1

[24] P. W. Emery, “Metabolic changes in malnutrition,” Eye, vol. 19, no. 10, pp. 1029–1034, Oct. 2005, doi: 10.1038/sj.eye.6701959.

[25] A. Eberstein and J. Goodgold, “Slow and fast twitch fibers in human skeletal muscle,” Am. J. Physiol.-Leg. Content, vol. 215, no. 3, pp. 535–541, Sept. 1968, doi: 10.1152/ajplegacy.1968.215.3.535.

[26] T. Jue, D. L. Rothman, B. A. Tavitian, and R. G. Shulman, “Natural-abundance 13C NMR study of glycogen repletion in human liver and muscle.,” Proc. Natl. Acad. Sci., vol. 86, no. 5, pp. 1439–1442, Mar. 1989, doi: 10.1073/pnas.86.5.1439.

[27] PubChem, “Cori cycle.” Accessed: Oct. 13, 2025. [Online]. Available: https://pubchem.ncbi.nlm.nih.gov/pathway/WikiPathways:WP1946

[28] P. B. Soeters, A. Shenkin, L. Sobotka, M. R. Soeters, P. W. de Leeuw, and R. R. Wolfe, “The anabolic role of the Warburg, Cori-cycle and Crabtree effects in health and disease,” Clin. Nutr. Edinb. Scotl., vol. 40, no. 5, pp. 2988–2998, May 2021, doi: 10.1016/j.clnu.2021.02.012.

[29] B. Bartoloni, M. Mannelli, T. Gamberi, and T. Fiaschi, “The Multiple Roles of Lactate in the Skeletal Muscle,” Cells, vol. 13, no. 14, p. 1177, July 2024, doi: 10.3390/cells13141177.

[30] M. Vanmunster et al., “Mechanosensors control skeletal muscle mass, molecular clocks, and metabolism,” Cell. Mol. Life Sci. CMLS, vol. 79, no. 6, p. 321, May 2022, doi: 10.1007/s00018-022-04346-7.

[31] B. S. Shenkman, “How Postural Muscle Senses Disuse? Early Signs and Signals,” Int. J. Mol. Sci., vol. 21, no. 14, p. 5037, July 2020, doi: 10.3390/ijms21145037.

[32] R. H. Westgaard and R. Bjørklund, “Generation of muscle tension additional to postural muscle load,” Ergonomics, vol. 30, no. 6, pp. 911–923, June 1987, doi: 10.1080/00140138708969787.

[33] S. K. Powers, M. P. Wiggs, J. A. Duarte, A. M. Zergeroglu, and H. A. Demirel, “Mitochondrial signaling contributes to disuse muscle atrophy,” Am. J. Physiol. - Endocrinol. Metab., vol. 303, no. 1, pp. E31–E39, July 2012, doi: 10.1152/ajpendo.00609.2011.

[34] K. Chen et al., “Forkhead Box O Signaling Pathway in Skeletal Muscle Atrophy,” Am. J. Pathol., vol. 192, no. 12, pp. 1648–1657, Dec. 2022, doi: 10.1016/j.ajpath.2022.09.003.

[35] J. Liang et al., “Lifelong Aerobic Exercise Alleviates Sarcopenia by Activating Autophagy and Inhibiting Protein Degradation via the AMPK/PGC-1α Signaling Pathway,” Metabolites, vol. 11, no. 5, p. 323, May 2021, doi: 10.3390/metabo11050323.

[36] R. Sartori, V. Romanello, and M. Sandri, “Mechanisms of muscle atrophy and hypertrophy: implications in health and disease,” Nat. Commun., vol. 12, no. 1, p. 330, Jan. 2021, doi: 10.1038/s41467-020-20123-1.

[37] J. Sheng et al., “LncRNA Foxo6os as a Novel ‘Scaffold’ Mediates MYBPC3 in Combating Pathological Cardiac Hypertrophy and Heart Failure,” Adv. Sci. Weinh. Baden-Wurtt. Ger., vol. 12, no. 34, p. e07365, Sept. 2025, doi: 10.1002/advs.202507365.

[S1] “Tissue fluid shift, forelimb loading, and tail tension in tail-suspended rats - NASA Technical Reports Server (NTRS).” Accessed: Oct. 27, 2025. [Online]. Available: https://ntrs.nasa.gov/citations/19850065036

[S2] R. K. Globus, D. D. Bikle, and E. Morey-Holton, “The Temporal Response of Bone to Unloading*,” Endocrinology, vol. 118, no. 2, pp. 733–742, Feb. 1986, doi: 10.1210/endo-118-2-733.

[S3] C. G. T. Tahimic et al., “Influence of Social Isolation During Prolonged Simulated Weightlessness by Hindlimb Unloading,” Front. Physiol., vol. 10, p. 1147, 2019, doi: 10.3389/fphys.2019.01147.

[S4] P. Di Tommaso, M. Chatzou, E. W. Floden, P. P. Barja, E. Palumbo, and C. Notredame, “Nextflow enables reproducible computational workflows,” Nat. Biotechnol., vol. 35, no. 4, pp. 316– 319, Apr. 2017, doi: 10.1038/nbt.3820.

[S5] P. A. Ewels et al., “The nf-core framework for community-curated bioinformatics pipelines,” Nat. Biotechnol., vol. 38, no. 3, pp. 276–278, Mar. 2020, doi: 10.1038/s41587-020-0439-x.

[S6] M. I. Love, W. Huber, and S. Anders, “Moderated estimation of fold change and dispersion for RNA-seq data with DESeq2,” Genome Biol., vol. 15, no. 12, p. 550, Dec. 2014, doi: 10.1186/s13059-014-0550-8.

[S7] A. Zhu, J. G. Ibrahim, and M. I. Love, “Heavy-tailed prior distributions for sequence count data: removing the noise and preserving large differences,” Bioinforma. Oxf. Engl., vol. 35, no. 12, pp. 2084–2092, June 2019, doi: 10.1093/bioinformatics/bty895.

[S8] V. K. Mootha et al., “PGC-1alpha-responsive genes involved in oxidative phosphorylation are coordinately downregulated in human diabetes,” Nat. Genet., vol. 34, no. 3, pp. 267–273, July 2003, doi: 10.1038/ng1180.

[S9] A. Subramanian et al., “Gene set enrichment analysis: A knowledge-based approach for interpreting genome-wide expression profiles,” Proc. Natl. Acad. Sci., vol. 102, no. 43, pp. 15545–15550, Oct. 2005, doi: 10.1073/pnas.0506580102.

