## Supplemental Figures 1-4 for "Fiber Type and Stimulus Determine Progression of Skeletal Muscle Atrophy"

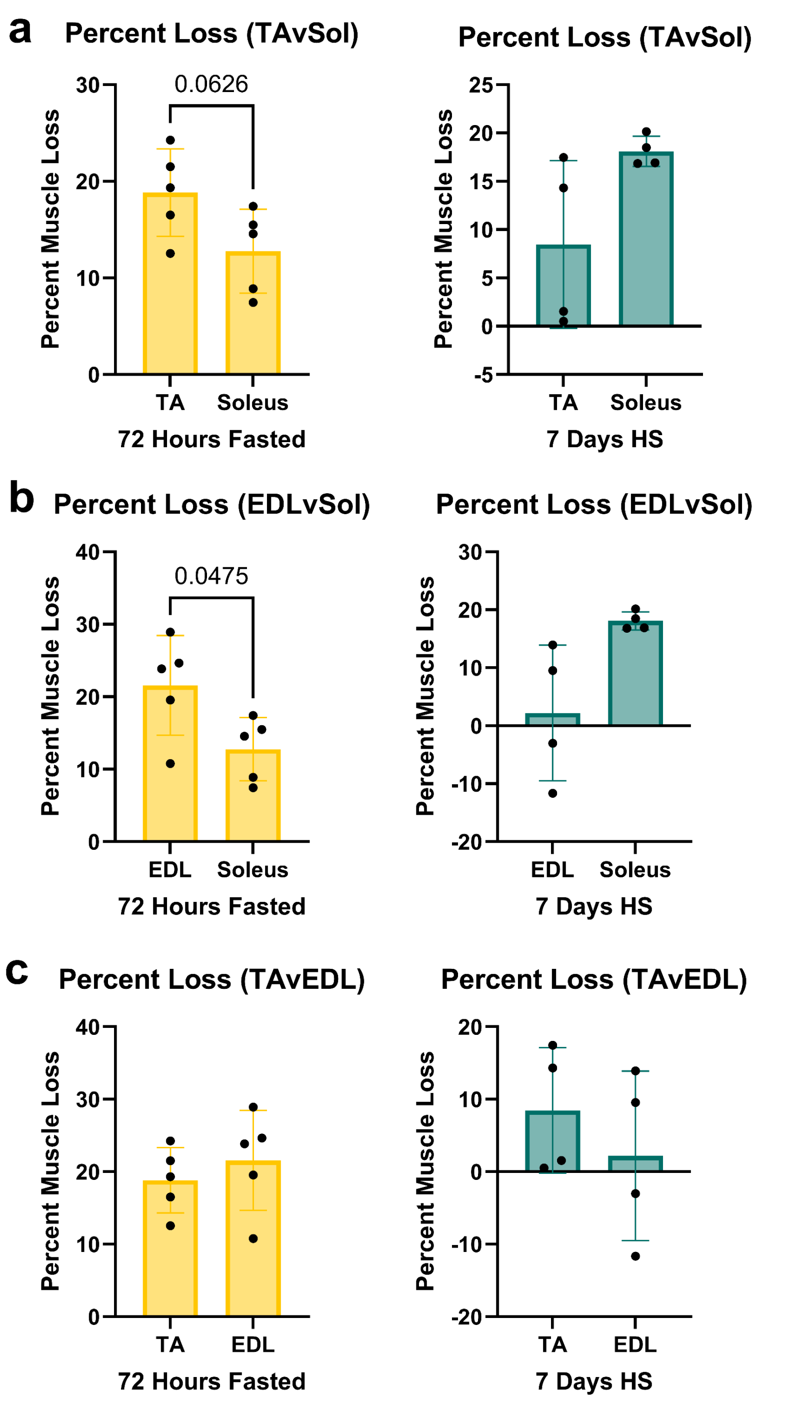


**Fig. S1 Fasting and hindlimb suspension cause variable muscle mass loss based on fiber type.** (a) Tibialis Anterior (TA) vs Soleus percent muscle loss after fasting (yellow, n=5) and hindlimb suspension (blue, n=4). (b) Extensor Digitorum Longus (EDL) versus Soleus percent muscle loss after fasting (yellow, n=5) and hindlimb suspension (blue, n=4). (c) TA vs EDL percent muscle loss after fasting (yellow, n=5) and hindlimb suspension (blue, n=4). All graphs were analyzed with Welch's t-test

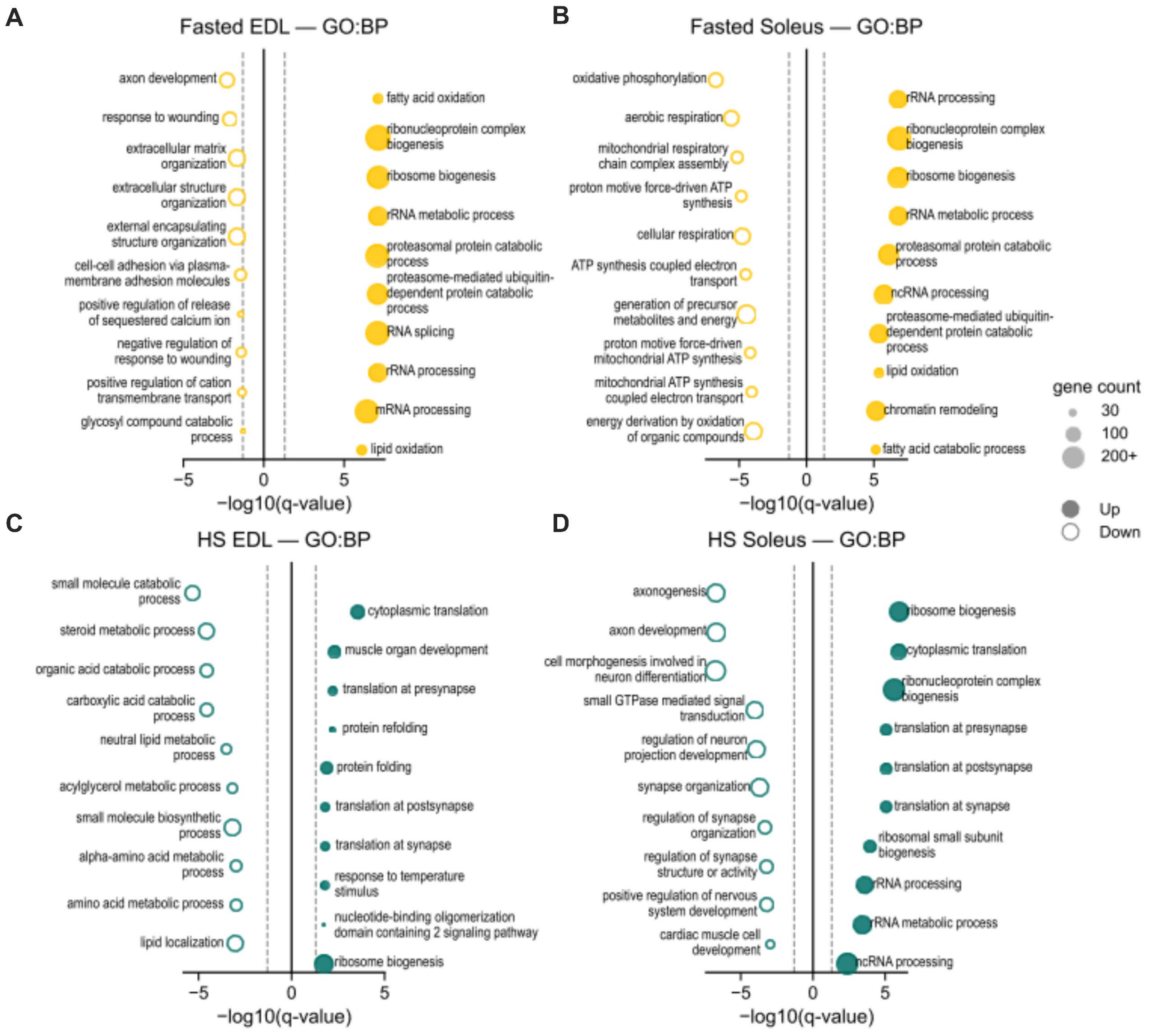


**Fig. S2 Enrichment analysis of significantly changing genes in the condition vs. control comparison for both EDL and soleus.** (a) GSEA Gene Ontology Biological Process for Soleus Fast vs. Soleus Control. (b) GSEA Gene Ontology Biological Process for EDL Fast vs. EDL Control. (c) GSEA Gene Ontology Biological Process for Soleus Suspension vs. Soleus Control. (d) GSEA Gene Ontology Biological Process for EDL Suspension vs. EDL Control. Gene set enrichment significance cutoff: q-value < 0.05; gene count = core enrichment genes.

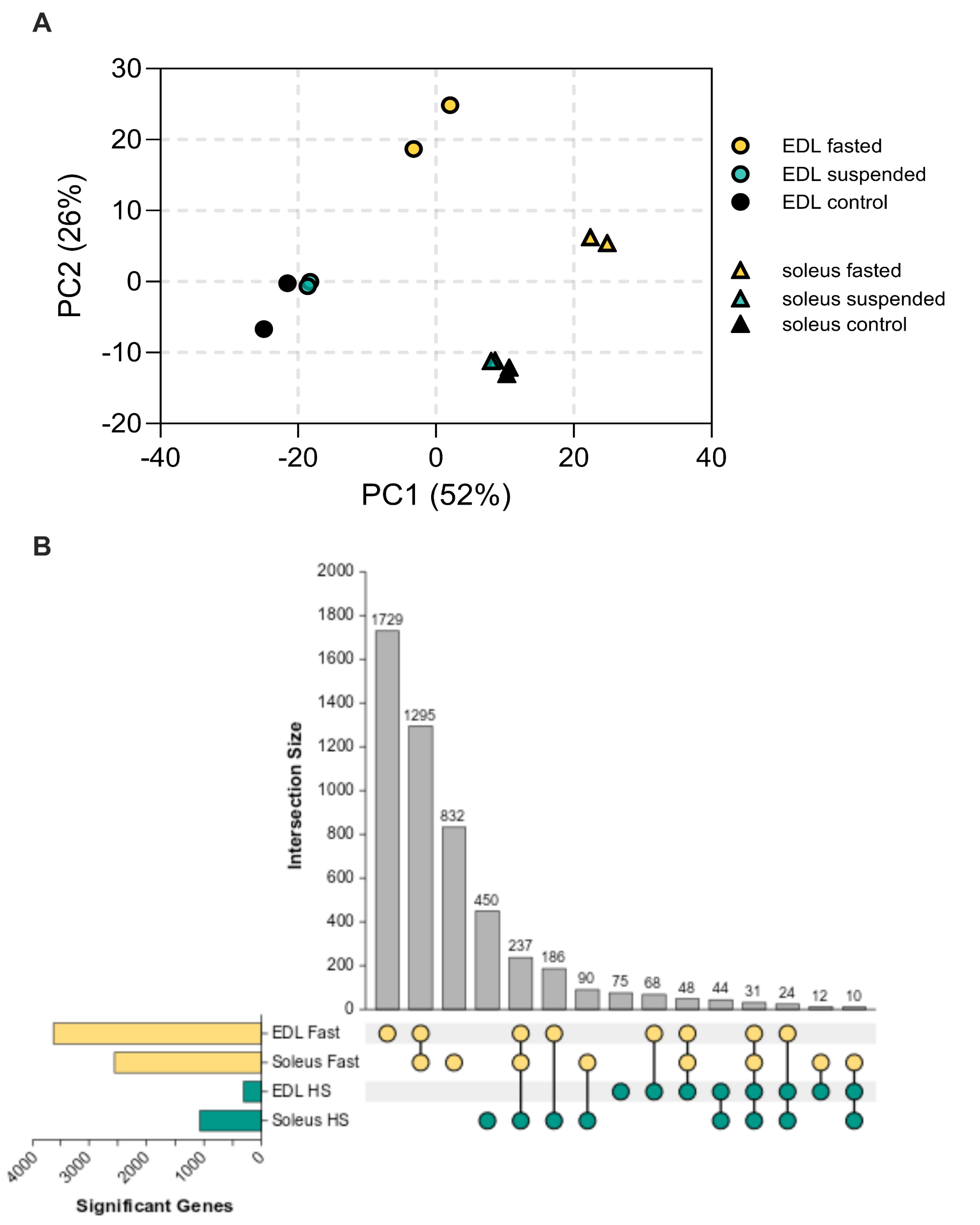


**Fig. S3 Dimensionality reduction and set overlap of transcriptional changes caused by fasting and disuse while accounting for muscle-type.** (a) Principal Component Analysis showing EDL (circles) and soleus (triangles) in either fasting (yellow), suspension (teal) or control (black) conditions (b) UpSet plot showing the set overlap of genes with significantly changing expression in each muscle-type and after each condition in a condition vs control comparison. Gene significance is determined by p-adj <0.05 and |fold-change| > 1.5.

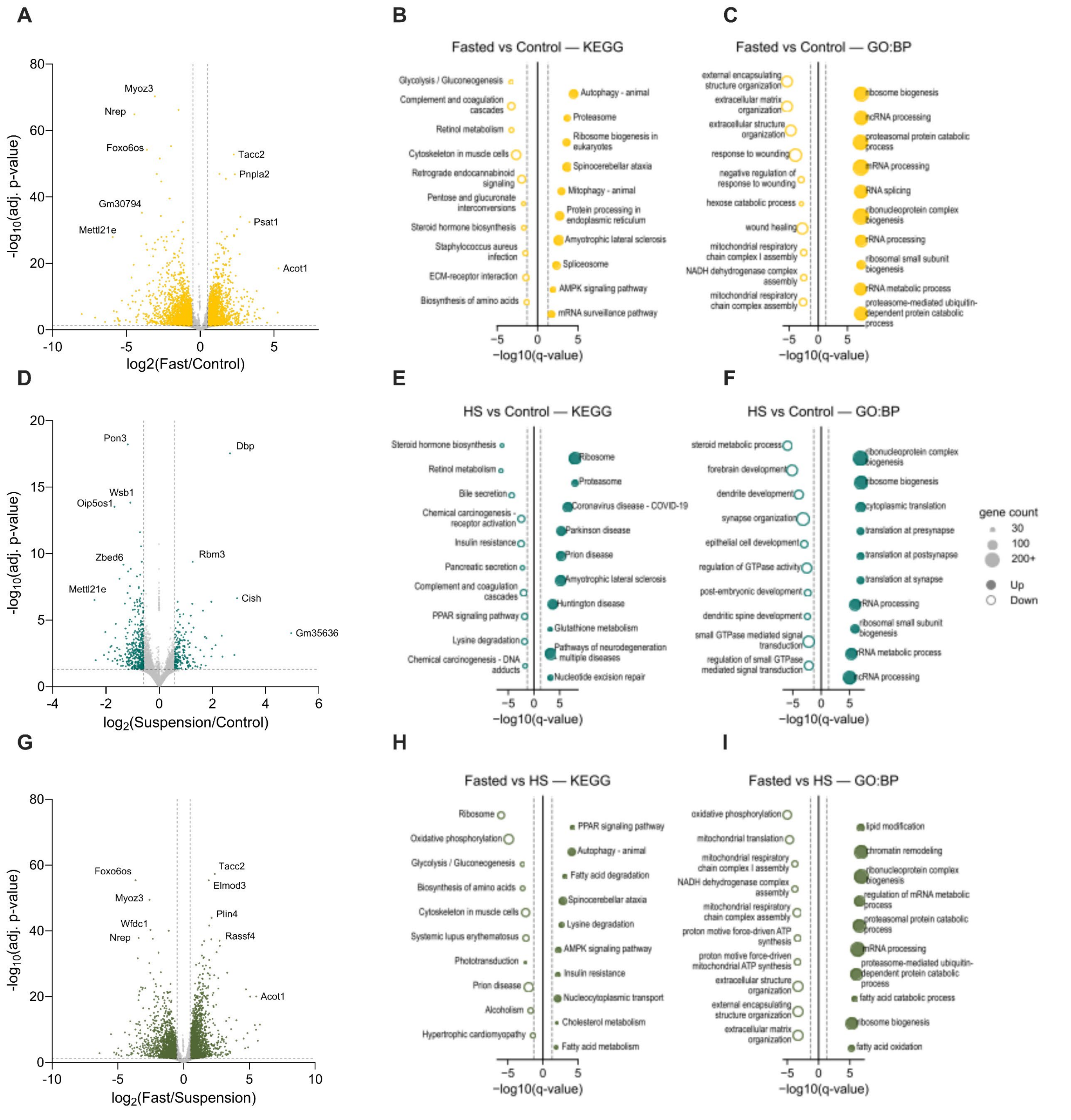


**Fig. S4 Disuse and nutrient deprivation elicit distinct transcriptional responses in skeletal muscle.** (a) Volcano plot for Fast vs. Control gene expression. (b-c) KEGG pathway (b) and GSEA Gene Ontology Biological Process (c) for differentially expressed genes in Fast vs. Control. (d) Volcano plot for Suspension vs. Control gene expression. (e-f) KEGG pathway (e) and GSEA Gene Ontology Biological Process (f) for differentially expressed genes in Suspension vs. Control. (g) Volcano plot for Fast vs. Suspension gene expression. (h-i) KEGG pathway (h) and GSEA Gene Ontology Biological Process (i) for differentially expressed genes in Fast vs. Suspension. Volcano plot significance cutoff = adj. p value < 0.05; fold change cutoff = |fold change| > 1.5; Gene set enrichment significance cutoff: q-value < 0.05; gene count = core enrichment genes
